# Directional Cell Migration Guided by a Strain Gradient

**DOI:** 10.1101/2021.07.07.451494

**Authors:** Feiyu Yang, Pengcheng Chen, Tianfa Xie, Yue Shao, Bo Li, Yubing Sun

## Abstract

Strain gradients, a graded change in the percentage of the deformation across a continuous field by applying forces, widely exist in development and physiological activities. The directional movement of cells is essential for proper cell localization, and directional cell migration in responses to gradients of chemicals, rigidity, and density and topography of extracellular matrices have been well-established. However, it is unclear whether strain gradients imposed on cells are sufficient to drive directional cell migration. In this work, we develop a programmable uniaxial cell stretch device coupled with geometrical constraints to create controllable strain gradients on cells. We demonstrate that single rat embryonic fibroblasts respond to very small strain gradients. In a gradient level of ∼4% per mm, over 60% of the REFs prefer to migrate towards the lower strain side in both the static and the 0.1 Hz cyclic stretch conditions. We confirm that such responses to strain gradient are distinct from durotaxis or haptotaxis. Moreover, we discover that the directional migration of the cells is initiated by increased focal adhesion contact areas and higher rate of protrusion formation on the lower strain side of the cell. We further establish a 2D extended motor-clutch model to explain the molecular mechanism. Through our model, we find that the strain-introduced traction force determines integrin fibronectin pairs’ catch-release dynamics, which drives such directional migration. Together, our results establish strain gradient as a novel cue to regulate directional cell migration and may provide new insights into development and tissue repairs.

## INTRODUCTION

Cells are constantly exposed to mechanical strains due to tissue growth (1), fluid flow (2), muscle contraction (3), etc. Numerous in vitro and in vivo studies support that mechanical strain is critically involved in various developmental and physiological processes, including the maturation of cardiac tissues, lung remodeling, and epithelial regeneration (4-5). While the majority of current research focuses on the cellular responses to uniform strains, strain gradients also widely exist due to heterogeneous tissue mechanical properties and bending/torsional moment (6-7). It has been observed in Xenopus ectoderm tissues that neuroepithelial cells collectively migrate along a strain gradient when subjected to concentrated loading, a process termed as “tensotaxis” (8). This phenomenon is drastically different from the extensively reported observation that cells reorientated under cyclic stretching (9-11). However, it is still unclear whether tensotaxis also regulates the directional migration of mesenchymal-like cells (12-13).

Directional cell migration can be guided by both biochemical and biomechanical cues in the cell microenvironment, including gradients of diffusible biomolecules (chemotaxis) (14), substrate bonded proteins (haptotaxis) (15, 16), substrate topography (topotaxis) (17), and the stiffness of extracellular matrix (durotaxis) (18, 19). Tensotaxis has yet been established as a biomechanical cue to guide directional cell migration, partially due to the difficulty to generate a controllable and physiologically relevant strain gradient (1-100% mm^-1^) (20), and more importantly, the challenge to distinguish durotaxis and tensotaxis, as substrate stiffness changes with strain due to nonlinear material responses (21, 22). Thus, tensotaxis and durotaxis are often used interchangeably, referring to the mechano-responsiveness of cells (23), while cells may utilize completely different mechanisms to sense strains and substrate stiffness.

Non-uniform strain fields have been generated using various approaches (24-28). In those works, it is consistently reported that cell reorientation is a function of strain magnitude and cells tend to avoid strain gradient and angle perpendicular to the principal strain directions. However, none of the existing systems produce a consistent strain gradient. As a result, the tensotaxis behaviors have not been observed. In this work, we developed a novel strain gradient generation device by introducing void regions with defined geometries to a membrane. Arduino microcontroller was used to precisely control the frequency and magnitude of uniaxial stretch with a servomotor. Using this device, we examined whether fibroblasts, which underwent constant stretches in vivo (29), respond to strain gradients directly. We further analyzed the role of adhesions and formation of protrusions in tensotaxis experimentally and computationally by developing an extended 2D motor clutch model.

## RESULTS

### Design and fabrication of the device for generating strain gradients

To study the effects of strain gradient on cell migration, it is essential to generate a controllable strain gradient to allow the migration of single cells in an optimal culture environment. Devices using microfluidic or vacuum to stretch cells can generate non-uniform strains. However, it is difficult to maintain a consistent strain field with a uniform gradient (30, 31). On the other hand, some systems require the encapsulation of cells in a sealed configuration (32). It is difficult to maintain the oxygen, pH, and nutrition conditions at optimal levels for long-term cell culture, and the profusion flow may introduce undesired shear stress, causing unexpected cellular responses (33).

To address these issues, we designed a new device for generating controllable strain gradients. This device was comprised of two primary components, the cell culture chamber and the control module (**Fig. 1a**). In the control module, a programmable servo motor attached with a rotational gear was fixed in the middle. Two translational gears were tightly jointed with the rotational gear (**Fig. 1b**). By controlling the rotational gear’s motion on the servo motor, the translational gears were driven to perform a linear movement for uniaxial cell stretching. The strain gradients were generated by a double-layer membrane mounted at the end of the translational gears (**Fig. 1c**). The bottom layer is a silicone film (1/32 inch) with cut-out of desirable geometries produced by laser cutting. On the top of the silicone base, we plasma-bonded a thin layer of polydimethylsiloxane (PDMS) membrane. Two glass slides were plasma-bonded at both ends of the silicone base (**Fig. 1c**). The advantage of this two-layer design is that the strain gradient can be modulated by the geometry of the cut-out, which is independent of the stretching magnitude and surface material properties. The assembly was mounted to the stretching device by clamping the two glass slides tightly with acrylic screws (**Fig. 1d**). During the experiments, cells were seeded on the PDMS membrane in the area with the triangular and squire cut-off. A 60 mm petri dish was placed underneath to submerge the cell into culture media. (**Fig. S1**).

**Fig. 1.**
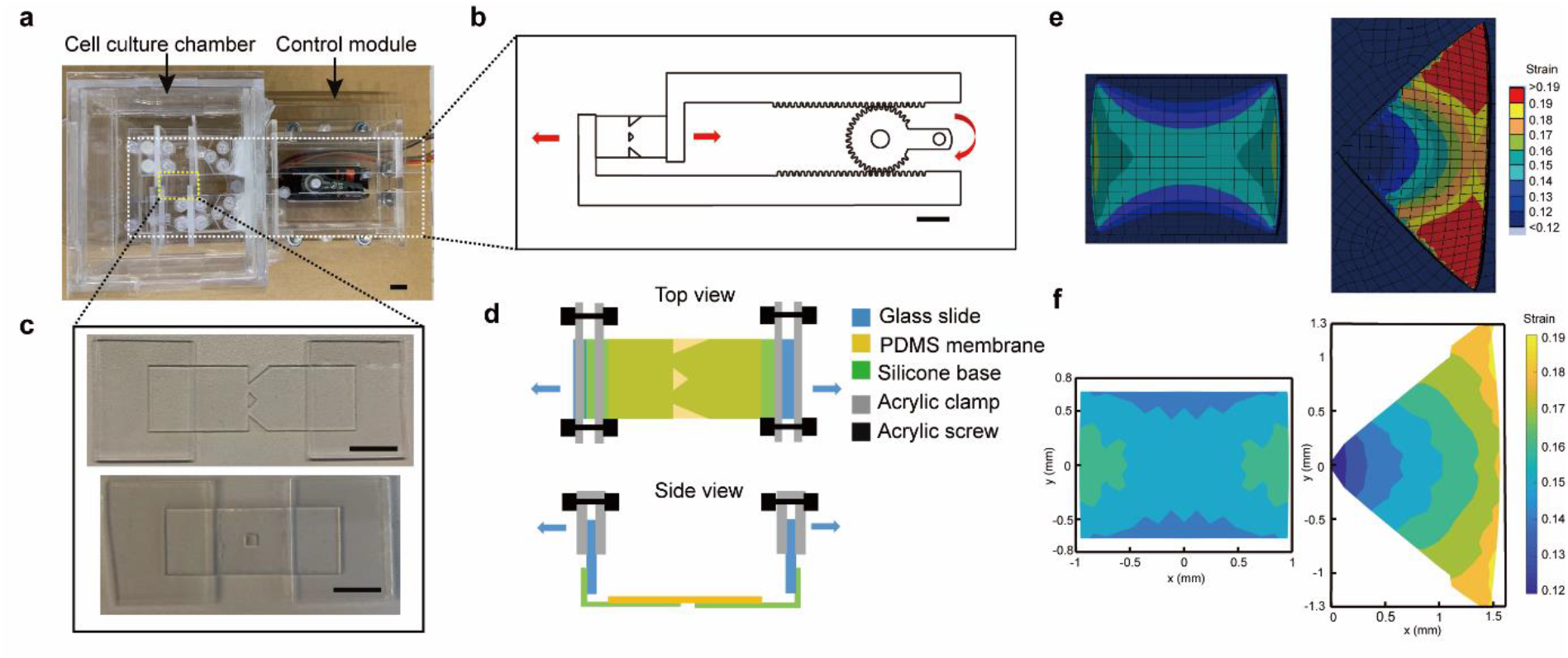
Design and calibration of the strain gradient generation device. **(a)** Photo of the strain gradient generation device. Scale bar, 10 mm. **(b)** Schematic showing the gear control mechanism. Scale bar, 10 mm. **(c)** Photo of double-layer membranes with triangle cut-out (top) and square cut-out (bottom). Scale bar, 10 mm. **(d)** Schematic showing device assembly and stretching application. **(e)** Simulated strain fields for uniform strain (left) and strain gradient (right). **(f)** Strain map showing experimental calibration of strain fields for uniform strain (left) and strain gradient (right).

### Characterization of the strain field

To establish the correlation between the cut-out geometry and the strain gradient, we first used finite element analysis (Ansys) to simulate the strain field under uniaxial stretching. A graded and a uniform strain field were established on the cell culture areas for the triangular and the square designs, respectively (**Fig. 1e**). To validate the simulation results, we further sought to characterize the device by mapping the strain field experimentally. we applied the micro-contact printing technique to print small markers with equal distance across the cell culture surface to calculate the strain across the cell culture area (**Fig. S2**). Images were taken before and after applying the stretch, and the displacement of markers was tracked using ImageJ (**Fig. S2a**). The strain and compression between two adjacent markers were calculated by dividing the change of distance by the initial distance in the x and y direction, respectively (**Fig. S2b**). Consistent with the simulation results, we found that stretching the membrane with triangular cut-out by applying 15-degree rotation with the rotational gear led to strains between 12% to 18% across a horizontal distance of 1.5 mm, equivalent to a strain gradient of ∼4% mm^-1^, while a uniform 15% strain was found for the membrane with square cut-out by applying 20-degree rotation with the rotational gear (**Fig. 1f, Fig. S3**). Notably, the compression generated due to stretching was negligible (**Fig. S4)**. As shown in **Fig. S3 and Fig. S4**, the sample-to-sample variations were small, suggesting this approach could reproducibly generate controllable strain gradients. The detailed experiment procedures and simulations can be found in the Method section.

It is possible that after extended cell culture or cyclic stretching, the changes of the material properties of the membrane may influence the strain field. To evaluate the stability of the strain gradient under these experimental conditions, samples were calibrated first, submerged into 1× DPBS in a 37 °C incubator for 24 hrs, cyclic stretched at 0.1 Hz for 3 hrs, and then were calibrated again. We found no significant changes in the strain field, suggesting that in our experimental conditions, the strain field remained stable.

### REFs migrate directionally towards lower strain direction

We next investigated whether the migration of single REFs could be influenced by the strain gradient. To avoid the haptotaxis effect caused by non-uniform ligand density due to the presence of strain gradient, we first stretched the membranes to desirable magnitudes, coated the membrane with fibronectin (FN) for 1 hr, and then released the membrane to the relaxed status. Single REFs were seeded on unstretched membranes with triangular and square cut-outs. After 15 hrs of culture, static stretching was applied to the membrane using the same conditions as in **Fig. 1f**, so that the gradient and uniform conditions have a comparable average strain of 15%. The membranes were held at the stretched status for the next 6 to 8 hrs and cell migration trajectories were tracked using live-cell microscopy (**Fig. 2a, Video S1-S5**). To avoid potential artifacts caused by membrane boundaries, only the cells in the center of the cell culture chamber were tracked. To minimize the influence of intercellular interactions, cells were seeded at a low density, and only samples with a total cell number of less than 170 cells were analyzed. Notably, only a few cells divided within the first 24 hours after cell seeding, and the cells that divided during the experiment period would not be tracked.

**Fig. 2.**
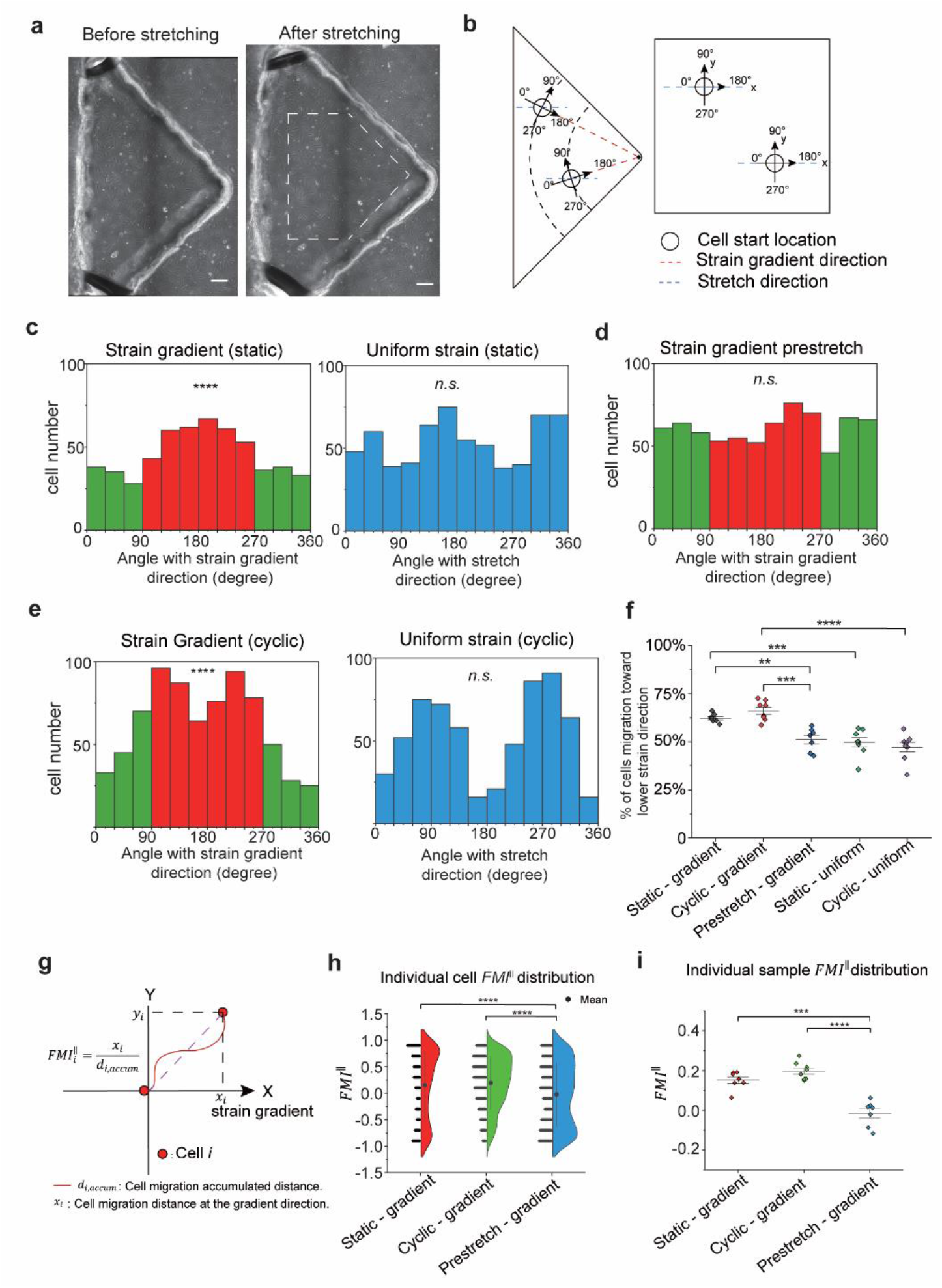
Tensotaxis behavior of REFs. **(a)** Photos of REFs seeding on the gradient double layer membrane, before (left) and after (right) stretching. Region of interests was encircled with white dash line. Scale bar, 200μm. **(b)** Schematics for the adjusted reference based on the cell’s location on the gradient (left) and the uniform double-layer membrane (right). **(c-e)** Histogram showing cell migration direction distributions under static strain gradient group (**c**, left, *N* = 7, *M* = 554), static uniform strain group (**c**, right, *N* = 8, *M* = 654), pre-stretched static strain gradient control group (**d**, *N* = 7, *M* = 732), cyclic strain gradient group (**e**, left, *N* = 7, *M* = 649), and cyclic uniform strain group (**e**, right, *N* = 8, *M* = 630). **(f)** Individual sample’s percentage of cell migrate to lower strain direction. Two sample t-tests were run between each pair of groups. **(g)** Schematic showing the *FMI*^||^ calculation. **(h)** Violin plots show individual cell’s *FMI*^||^ distribution for static gradient group (*N* = 7, *M* = 554) (Mean =0.158), cyclic gradient group (*N* = 7, *M* = 649) (Mean= 0.228), and pre-stretched strain gradient control group (N = 7, *M* = 732) (Mean=-0.022). Mann-Whitney tests were run between each pair of groups. **(i)** Individual samples’ average *FMI*^||^ distribution for static gradient group (*N* = 7), cyclic gradient group (*N* = 7), and pre-stretched strain gradient control group (*N* = 7). Two sample t-tests were run between each pair of groups. Rayleigh tests were run to determine the unimodal distribution of the circular data. **, *P* < 0.01. ***, *P* < 0.001. ****, *P* < 0.0001. *N*: Sample number; *M*: Cell number.

To quantify the directionality of the cell migration relative to the strain gradient direction, we set up local coordinates for each cell with x-axis being the maximum gradient direction (**Fig. 2b**). We then connected the first and the last cell coordinate during the cell tracking period to calculate the migration angles. The cells were considered migrating towards the lower strain direction when the migration angle was between 90º to 270º. On the other hand, when the migration angle was between 0º to 90º or 270º to 360º, the cells were considered migrating towards the higher strain direction (**Fig. 2b**). For the uniform strain conditions, as no strain gradient was established, the x-axis was set to be along the stretch direction.

Histograms were plotted for visualizing cell migration direction preferences (**Fig. 2c**). We found 62.45% of cells migrating towards the lower strain direction in the static gradient group, which was significantly larger than the higher strain direction as confirmed by the Rayleigh test (**Fig. 2c**). In contrast, in the uniform strain condition, no preference in the directionality of cell migration was found **(Fig. 2c)**.

### Tensotaxis is distinct from durotaxis

When the PDMS membranes were stretched, the strain gradient could potentially lead to a variation of stiffness across the PDMS membrane due to the non-linear mechanical property of PDMS (34, 35). Therefore, the directional cell migration we observed might be induced at least partially by durotaxis. Moreover, micro-size fibrillar structures could be established on the PDMS membrane due to stretching, which might affect cell migration preferences (36). To exclude the influence of these factors, we design a pre-stretched static group as a control (**Fig. 2d**). We first stretched the membrane and then coated it with FN to create a uniform ECM coating. We next seeded cells and tracked the cell migrations after 15 hrs of incubation. In this way, cells would not be subjected to stretch, therefore not being exposed to the strain and the strain gradient, while the condition of the substrates was the same as the static strain gradient group shown in **Fig. 2c**. No significant preference of cell migration was found in this pre-stretched condition, suggesting that cells directly sense the strain gradient imposed on cell bodies, rather than substrate material properties.

### Cyclic stretching induces cell migration perpendicular to stretching direction

We next investigated whether cyclic stretching induced different migration patterns. Both membranes with triangular and square cut-out were cyclically stretched for the first 3 hours at Hz, then were held at the stretched status for the next 3 hours. Interestingly, more cells migrated to the direction perpendicular to the strain gradient / stretching directions (migration angle close to 90º and 270º), compared to the static condition (**Fig. 2e**). This trend is more prominent for the uniform strain condition. However, under cyclic stretching conditions, we still observed a significantly high percentage of cells (66.35%) migrated towards the lower strain direction for the strain gradient condition (**Fig. 2e**). These data suggested that cell reorientation to avoid strain gradients and tensotaxis are two independent, competing mechanisms to regulate directional cell migration upon stretching. This observation is further confirmed by the statistical analysis showing that biased cell migration towards the lower strain direction only existed in the presence of a strain gradient (**Fig. 2f**).

### Quantifying tensotaxis using the forward migration index

We next evaluated the efficacy of the tensotaxis by quantifying the forward migration index (*FMI*^||^) (37, 38), the ratio of the cell migration distance in the maximum gradient direction to the accumulated distance (**Fig. 2g**). When the *FMI*^||^ equals to 1 or -1, the cell migrates along or against the maximum gradient direction with no deviations, respectively. As such, a higher value of *FMI*^||^ represents a higher efficacy of tensotaxis response. Consistent with the histograms, we found that both static and cyclic stretching of cell-seeded samples with strain gradient led to more cells with the *FMI*^||^ closer to 1, with an average *FMI*^||^ of 0.158 and 0.228, respectively (**Fig. 2h-i**). In comparison, the pre-stretched condition led to an average *FMI*^||^ closer to 0 (**Fig. 2h-i**). Notably, the *FMI*^||^ for durotaxis of multiple cell types are smaller than 0.2 (39), suggesting that strain gradient is a potent cue to drive directional cell migration.

### Strain magnitude regulates tensotaxis response rate

We next investigated how strain magnitude regulated the tensotaxis. We divided each sample’s cell culture area into four regions (*R1* to *R4*) based on the strain magnitude (13.5% - 15%, 15% - 16%, 16% - 17% and 17%-18%) with a consistent strain gradient of about 4% mm^-1^. (**Fig. 3a**). We compared the *FMI*^||^ for cells in each region under static gradient, cyclic gradient, and pre-stretch conditions. As shown in **Fig. 3b-c**, we found that in both static and cyclic gradient groups, the *FMI*^||^ was significantly larger for the cells in *R4*. We also observed trends of average *FMI*^||^ increasing from *R1* to *R4* for the static and cyclic groups (**Fig. 3e)**, suggesting a higher strain magnitude triggers a stronger tensotaxis response. No significant differences were found in the pre-stretched group. Together, these results suggest that a higher strain magnitude can increase the tensotaxis response rate.

**Fig. 3.**
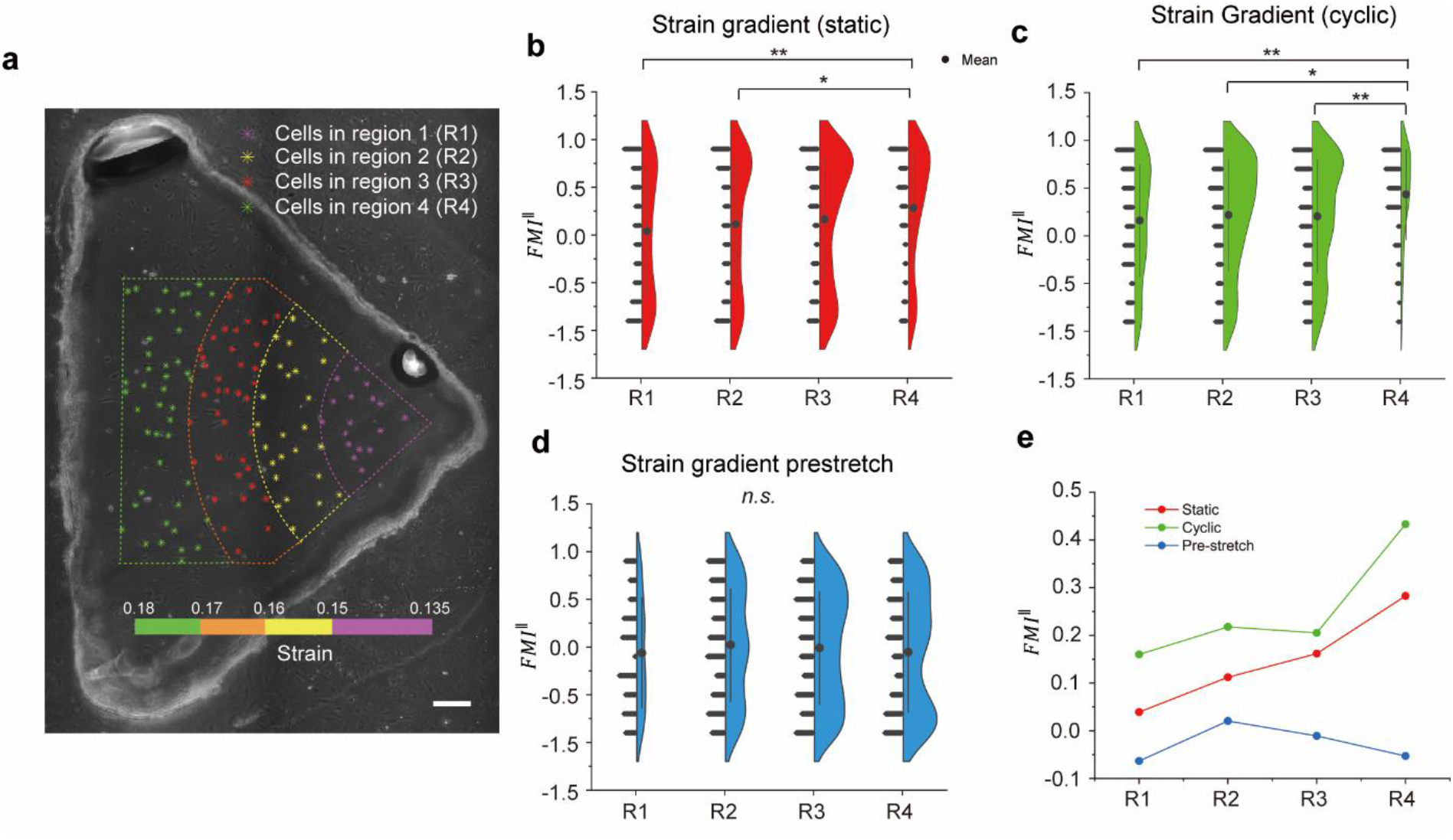
Strain magnitude regulates REF tensotaxis. **(a)** Photo of dividing an individual gradient sample into four strain magnitude regions (*R1* to *R4*). Scale bar: 200 μm. (**b-d**) Violin plots show individual cell’s *FMI*^||^ distribution from *R1* to *R4* for the static gradient group (**b**, *N* = 7, *M1* = 94, *M2* = 128, *M3* =201, *M4* =131), cyclic gradient group (**c**, *N* = 7, *M1* =116, *M2* = 242, *M3* =223, *M4* =68), and pre-stretched static gradient control group (**d**, *N* = 7, *M1* = 66, *M2* = 171, *M3* =247, *M4* = 248). (**e**) Individual cells’ average *FMI*^||^ from *R1* to *R4* for the static gradient, cyclic gradient and pre-stretched control group. Mann-Whitney tests were run between each pair of groups. *, *P* < 0.05, **, *P* < 0.01. *N*: Sample number; *M1*: Cell quantity in *R1*; *M2*: Cell quantity in *R2*; *M3*: Cell quantity in *R3*; *M4*: Cell quantity in *R4*.

### Increased focal adhesion formation and cell protrusion on the lower strain side of cells

We next investigated molecular mechanisms for the tensotaxis. As actin polymerization-driven cell protrusion and the formation of new adhesion sites are the two most critical steps in cell locomotion (40), we sought to investigate the directionality of single-cell protrusion and focal adhesion dynamics under a strain gradient. The REF52 cells have been previously transfected with a YFP-Paxillin reporter (41), and live-cell microscopy was used to evaluate the dynamics of paxillin containing focal adhesions. Cells cultured on membranes with triangular cut-out were imaged before and 20 mins after stretching (**Fig. 4a**).

**Fig. 4.**
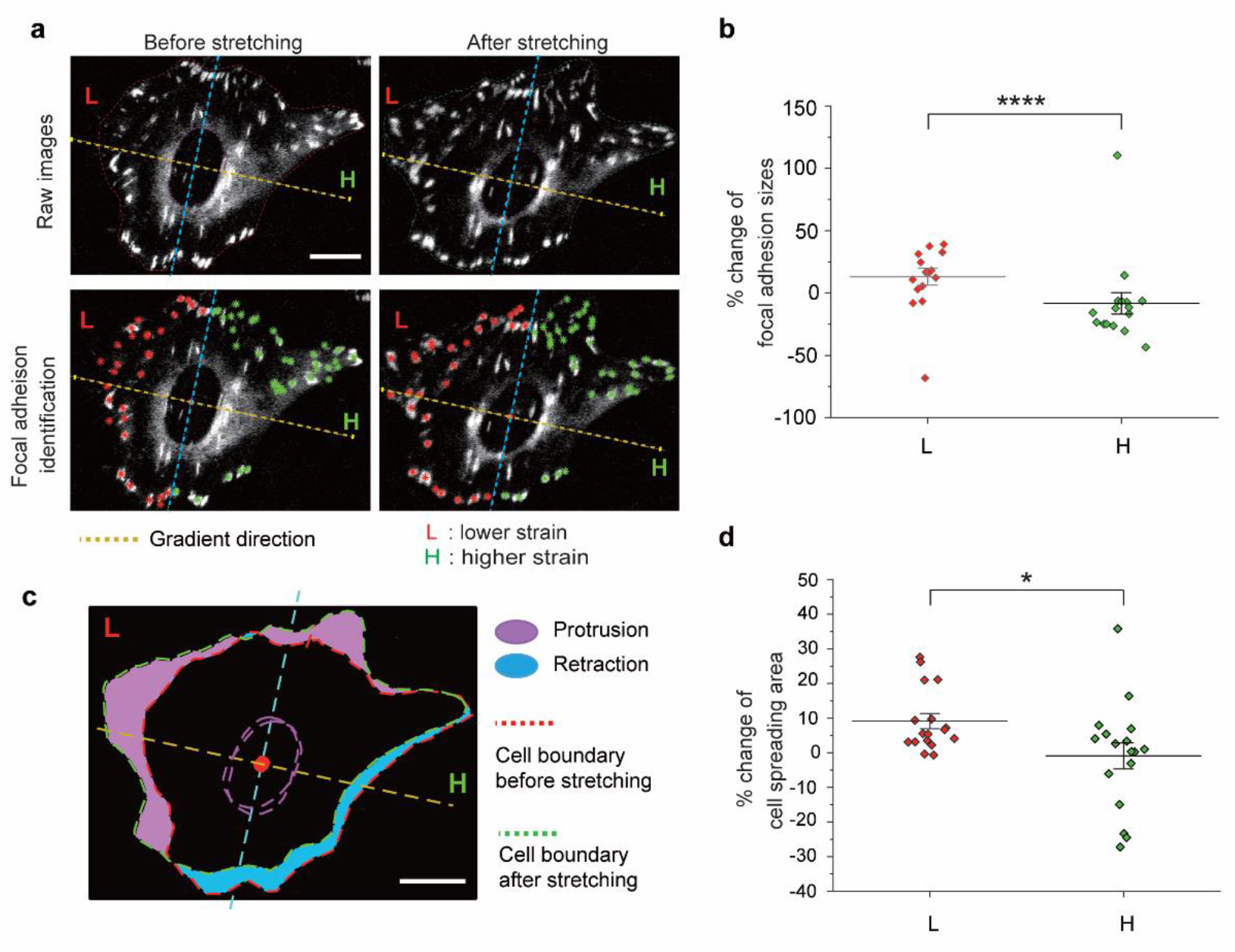
Focal adhesion dynamics and polarized cell protrusion in response to strain gradient. **(a)** Raw images (top) and focal adhesions identified images (bottom) of a representative REF before (left) and after (right) stretching. Scale bar 20 μm **(b)** Normalized percentage change of the total focal adhesion area in the lower and higher strain halves of the cells (*N* = 16). **(c)** Overlapping the images before and after stretching by the nucleus center (red dot) and the gradient direction (yellow dash line). The protrusions (purple) and retractions (blue) after stretching were color coded. **(d)** Percentage of cell area change in the lower and higher strain half (*N* = 17). Two sample t-test were run. *, *P* < 0.05. ****, *P* < 0.0001. *N*: cell number.

To analyze the focal adhesion dynamics as a function of strain gradient, we divided each cell into two halves along the axis passing the center of the cell nucleus and perpendicular to the strain gradient direction (blue line in **Fig. 4a**). We used ImageJ to automatically identify each focal adhesion and quantified their sizes. After compensating for the imaging artifacts caused by stretching (see Methods for details), we found that the total focal adhesion area increased significantly on the lower strain half compared with the higher strain half after stretching (**Fig. 4b**).

We next quantified the cell protrusion and retraction on the lower and higher strain sides. We found that within 20 mins after stretching, a significant protrusion on the lower strain half of the cell and retraction on the higher strain half could be found in the majority of cells analyzed (**Fig. 4c-d**). By simultaneously analyzing focal adhesion dynamics and cell protrusion, we found that 62.5% of cells have both a higher rate of protrusion formation and a relative increase of focal adhesion contact area at the lower strain direction (*n* = 16), which conformed to the tensotaxis response rate shown in **Fig. 2c**. Together, these results suggest that protrusion and preferential formation of focal adhesions on the lower strain side of the cell may lead to the tensotaxis of cells.

### Extended Motor-clutch model recapitulates tensotaxis

To explain the mechanism of the tensotaxis behavior governed by focal adhesion dynamics, we developed a modified version of the extended 2D motor-clutch model (EMM) (42-43). We propose that the cell motility is determined by the catch-release dynamics of integrin and FN pairs, which are partially governed by strain gradient dependent cell traction forces. Here, an individual REF is modeled as a polygon attached to a stretched elastic substrate (**Fig. 5a)**, whose vertexes are connected by springs. The actin filaments connect the cell vertexes to the nucleus centroid (**Fig. 5b**). By incorporating the EMM with each filament to quantify the on-rate of integrin-FN bonds (*k*_*on*_), we found that the higher affinity of bonds appears on the lower strain side of the substrate, which is consistent with the focal adhesion dynamics observed in the experiment. We then simulated the migration of an individual REF coupled with 34 vertexes on a 4% strain gradient substrate for 6 hrs. We confirm that the EMM is sufficient to recapitulate the directional cell migrations (**Fig. 5c, d)**.

**Fig. 5.**
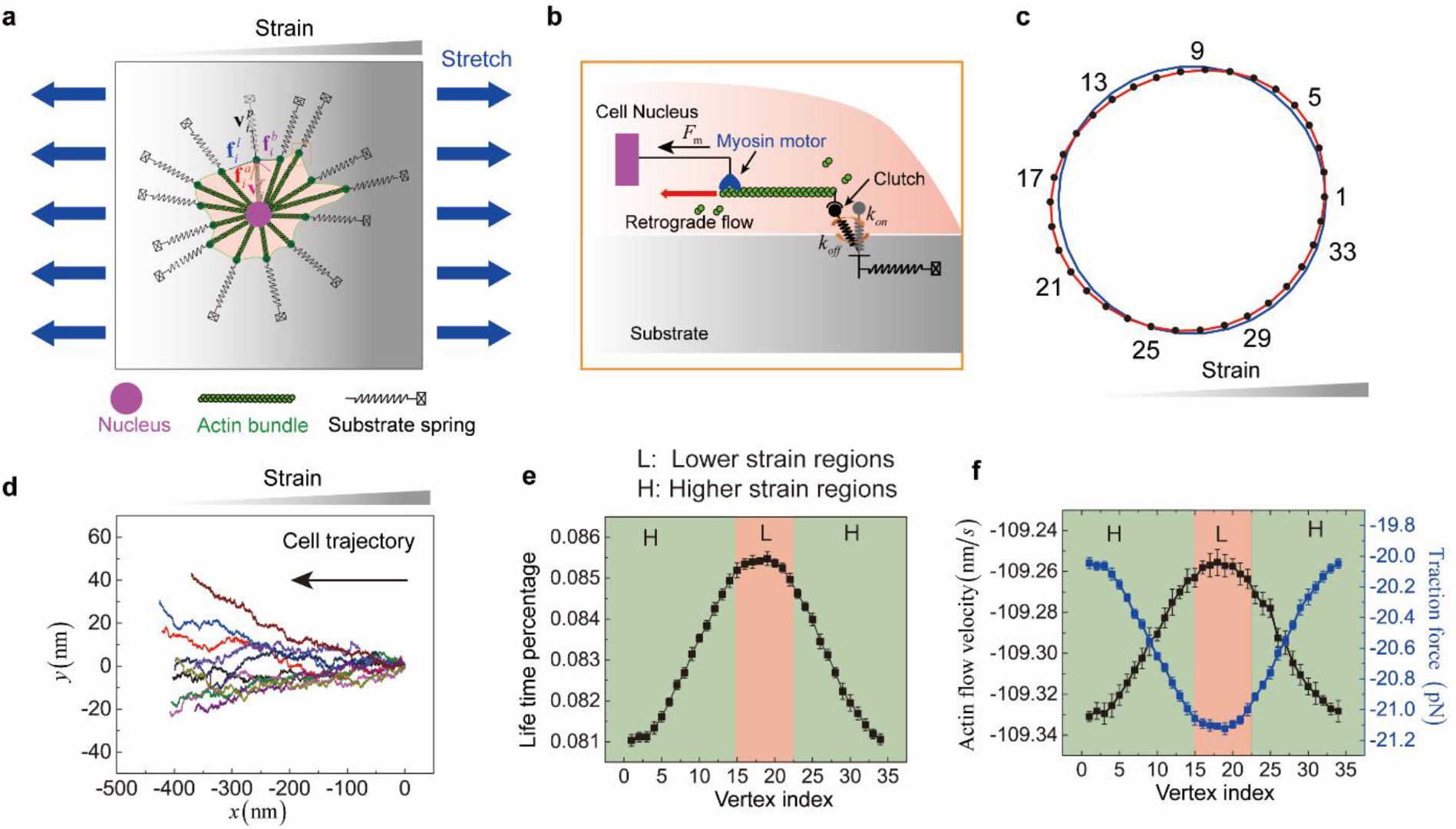
Extended 2D motor-clutch modeling simulates tensotaxis. **(a)** Schematic of a cell model. **(b)** Motor-clutch module of an individual vertex. **(c)** Schematic of a simulated cell model migrating from (blue line) to (red line) on a 4% gradient of strain substrate. The cell centroid is located on the 15% strain region. Black dots are cell vertex. The numbers indicate the vertex index. **(d)** Representative of predicted cell trajectories on the 4% gradient of strain substrate (*N*=10). Cell centroid (x=0, y=0) is located on 15% strain region. **(e)** Life time percentage of engaged integrin-fibronectin pairs **(f)** and traction force, and actin retrogradation flow velocity during cell migration. Red and green regions represent the lower and high strain side, respectively. *N*: cell number.

We further verified the consistency between EMM and experiment results by calculating the lifetime percentage *α* = α*t T*, where α*t* is the total time of engaging for integrin-FN pairs and *T* is the whole simulation time. As shown in **Fig. 5e**, longer lifetime of integrin-FN bonds is observed on the lower strain regions, resulting in focal adhesions with longer life spans and larger sizes. This result suggests higher traction and slower retrogradation speed of actin filament on the side with lower strain **(Fig. 5f**), which enables cell protrusion towards lower strain region. Together, our extended motor-clutch model can well recapitulate cell tensotaxis behaviors observed in our experiments.

## DISCUSSION

Many tissues with anisotropic mechanical properties were subject to constant deformation during development and normal physiological activities. While the cell reorientation under strain and strain gradient has been well-studied, it is unclear whether a strain gradient also guides directional cell migration. This is particularly important to fully understand the principles of morphogenesis in development. In this work, we show that fibroblasts are extremely sensitive to a small strain gradient (∼4% mm^-1^) and migrate toward the lower strain direction under static stretching. While cyclic stretching induced cell reorientation towards the direction perpendicular to cell stretching, we still observed an independent tendency for cell migration along the cell stretching direction toward the lower strain side. Our results demonstrate that the tensotaxis of cells depends on the magnitude of strain and is initiated by strain dependent focal adhesion formation and cell protrusion. Using a modified motor-clutch model, we successfully recapitulate the tensotaxis of cells by predicting the catch-release of integrin and FN bonds. Together, our data unambiguously establish the tensotaxis phenomenon of fibroblasts, which is distinct from durotaxis and haptotaxis.

The observation of the tensotaxis is allowed by our novel strain gradient generation device. Our approach allows a continuous strain gradient in mm scale for long-term cell migration tracking, distinct from the non-uniform strain field generated by applying concentrated forces (25). As the strain gradient can be tuned by changing the geometry of the cut-out in the underlying silicone layer without altering stretching parameters, the strain gradient and stretching magnitude, frequency, strain rate, and surface material properties can be controlled independently. No shear forces caused by the flow of media or air were introduced into the system. Compared to microfluidic-based and commercially available cell stretchers, our device is facile to fabricate without the need for cleanroom facilities and completely biocompatible, making it suitable for broader adaptations. Future works shall focus on modifying the device to study how strain gradient and other strain parameters synergistically regulate cell migration and other cell functions such as stem cell differentiation.

Our experiment (**Fig. 2d**) clearly demonstrated that tensotaxis is distinct from durotaxis and haptotaxis. No preference in cell migration directionality was found when the membrane was pre-stretched and coated uniformly with FN. This is likely due to the narrow range of substrate stiffness cells can sense (*E* = 10^−1^-10^2^ kPa), and that durotaxis was generally investigated using soft hydrogel (*E* = 2 – 7 kPa) (18). In contrast, the Young’s modulus for PDMS membrane was reported to be over 1MPa, on which durotaxis is not prominent (35, 40). It has been established that cells utilize mechanosensitive focal adhesions (44-45), filopodia structures (46), and contractile machinery (47) in their rigidity sensing. Our results in **Fig. 4&5** also show that the focal adhesions are sensitive to strain and the stability and/or formation of new focal adhesions are preferred in regions with lower strain, which may facilitate the protrusion of cells towards lower strain direction. One possible explanation is the catch-slip bond-like behaviors observed in focal adhesions (41). Smaller strain may lead to a force-dependent stabilization of focal adhesions (“catch”), while larger strain may lead to the dissociation of focal adhesions (slip). As such, we have confirmed the hypothesis through a modified version of the classic motor-clutch model, demonstrating that the strain gradient would control the binding and unbinding of integrin-FN bonds resulting in directional migration.

In conclusion, by generating a controllable strain gradient on cells, we demonstrated cells directly respond to a strain gradient and migrate directionally to the direction with lower strain. This mechanosensitive behavior, termed tensotaxis, is distinct from durotaxis or haptotaxis, and depends on the magnitude of the strain applied to the cells. Subcellular analysis revealed that lower strain increases the levels of focal adhesions and facilitates cell spreading. Simulations suggest that gradient-induced traction variation would determine the binding and unbinding of integrin-FN bonds, which drive the tensotaxis of cells. Together, we establish the strain gradient as a mechanical cue to guide the directional migration of single cells and provide insights into the mechanisms of tissue remodeling and morphogenesis.

## METHODS

### Device and double-layer membrane fabrication

Parts of the cell stretching device were printed with a laser cutter (40 Watt Epilog Mini 18 × 12) using the acrylic sheet (**Fig. 1a**), and assembled with screws and sealed with PDMS. In the control module, a programmable servomotor (DS3218, Annimos) attached with a rotational gear was fixed in the middle. The Arduino Uno microcontroller was adopted to control the servomotor. Two translational gears were properly joined with the rotational gear on two sides, which were designed to connect the cell culture chamber with the control panel (**Fig. 1b**). Two small openings in between allowed the translational gears to extend into the cell culture chamber. Parafilms (PM-996, Bemis) were wrapped around the translational gears to seal the openings (**Fig. S1**).

The cell culture chamber was encapsulated to create a contamination-free, biocompatible environment. The chamber could be opened from the top and the bottom (**Fig. S1**). We designed a removable lid to cover the top of the chamber, and small gaps were left to access air. On the bottom, an opening was made below the double-layer membrane, which was sealed with a removable acrylic sheet by screws.

The double-layer membrane was fabricated with 1/32 in silicone film, PDMS membrane, plasma bonded on two glass slides at both ends (**Fig. 1c, Fig. S5a**). The silicone film (base layer) was cut through with a laser cutter to create desired shapes. Dow Corning Sylgard 184 silicone elastomer and cure agent (GMID: 04019862) were mixed in the 10:1 ratio to fabricate the PDMS membrane (top layer). A small amount (<500 μl) of PDMS was pipetted on the center of an Ø85 mm acrylic circle. The PDMS was spin-coated the acrylic circle at 500 RPM for 30 s and then at 1000 RPM for another 2 mins (WS650MZ23NPPB, Laurell Technologies). PDMS-coated acrylic circles were cured in a 65 °C oven overnight. A thin layer of PDMS membrane with a thickness of around 100 μm was formed on top of the acrylic circle. The thickness was measured with a precise micrometer (293-340330, Mitutoyo). The PDMS coated acrylic was laser cut into the 22×14 mm and 14×14 mm pieces for the triangular and the squire cut-off, respectively (**Fig. 1c**). The glass slide (125444, Fisher Scientific) was cut into 20×25 mm pieces. We laser printed acrylic template in the shape of the double-layer membrane for precise alignment (**Fig. S5**). All parts were cleaned with 100% ethanol. The silicone film base, PDMS membrane, and glass slides were bonded with a plasma cleaner (PDC-001, Harrick plasma) at 500 psi for 3 mins, then baked at 65 °C overnight for firm bonding. The dimensions of the designs were illustrated in **Fig. S6**.

### REF cell culture in the cell stretching device

REF-52 expressing YFP-paxillin fusion protein (a gift from Dr. Jianping Fu) were cultured in T-25 flasks and subcultured at 90% confluency. The culture media were composed of DMEM (11960051, Gibco) supplemented with 10% FBS (10082147, Gibco), 1% MEM NEAA (11140050, Gibco), 1% GlutaMAX (35050061, Gibco), and 1% Penicillin Streptomycin (15140122, Gibco).

To prepare the device for cell seeding. The double-layer membrane was first sonicated for 5 mins in 100% ethanol to remove all particles on the surface, and then sterilized by autoclaving. The cell stretch device was sprayed with 70% ethanol, then blow-dried in a biosafety cabinet. To avoid the effect of haptotaxis, we stretched the membranes to the desired magnitudes before coating FN. The membranes were incubated at room temperature with 50 μg/ml FN (33016-015, Gibco) for one hour by pipetting 100 ml solution on the cell culture area, which was then rinsed off with 1× DPBS.

REFs were seeded at the density of 50 K to 80 K/ml by pipetting 100 μl cell suspensions on top of the cell culture chamber. The device was then placed into a 37 °C incubator for 1 hr for cell attachment, and then 5 ml culture media were added to submerge the cells for overnight incubation. The stretching and imaging procedures would begin 15 hrs after cell seeding.

### Finite Element Analysis

Finite element modeling of the double-layer membrane was conducted using the Ansys simulation software. Two-layer 3-D models were constructed to mimic the structures of the double-layer membrane (**Fig. S7a**). The PDMS membrane was defined to bond on top of the silicone base (**Fig. S7b**). For simplicity, only the cell seeding chamber was modeled. For the silicone film, a tensile test was conducted to acquire the test stress-strain data (Mark-10 Force Gauge Model M7-5), which was curve-fitted using the polynomial 2^nd^ order equation in the simulation (**Fig. S7c**). For the PDMS membrane, the material properties were defined based on previous literature (48), with Young’s modulus set to 1.1 MPa, the Poisson’s ratio to 0.45, and density to 970 kg/m^3^.

In the simulation, the forces applied in the x-direction were estimated to conform to the experimental strain magnitude across the cell culture area. To mimic the uniaxial stretching, forces were applied on both sides of the models (**Fig. S7a**). The parameters of the simulation were described in **Table S2**.

### Strain field calibration

To quantify the strain across the cell culture area, patterns of small markers (Ø 50 μm) with equal distance were transferred from microfabricated PDMS stamps using the micro-contact printing technique (49). Patterns with center to center distances of 80 μm and 100 μm were used. PDMS stamps were incubated with 50 μg/ml Alexa Fluor 555-labeled BSA (A34786, Molecular Probes) at room temperature for 1 hr, rinsed with DI water and blow-dried. The double-layer membranes were UV treated for 7 mins for surface activation (Model 30, Jelight Company). The fluorescence markers were printed by pressing the PDMS stamp onto the membrane.

The strain and compression were calculated by quantifying the change of distance between two adjacent markers along and perpendicular to the stretching direction, respectively (**Fig. S2**). The strain/ compression fields were mapped based on the coordinates of the markers after stretching and the corresponding strain/compression using the Matlab *contour* (**Fig. S3-4)**. The strain/compression fields were symmetrical based on the x-axis starting from the triangle’s vertex for the triangular design in the stretch direction. For the square sample, the strain/compression field was symmetrical both in the x and y directions to the sample’s center. To reduce variations during calibration, the strain/compression fields were averaged based on the respective symmetry axes for both designs.

Three samples for both designs were calibrated (**Fig. S3, 4**). To compare each sample’s strain/compression fields, we plotted the strain/compression magnitude against the distance from the origin in the direction L1, L2, and L3 (**Fig. S3b, d, Fig. S4b, d**). For the triangular design, the origin was the vertex (**Fig. S3a, Fig. S4a**), and the L1 was the line starting from the vertex to the x-direction. L2 and L3 were acquired by rotating L1 for 15 and 30 degrees counterclockwise, respectively. For the square design, the origin was the square center (**Fig. S3c, Fig. S4c**), L1 started from the center to the x-direction, and L2 and L3 were acquired by rotating L1 for 45 and 90 degrees counterclockwise, respectively. Data points close to the reference lines were used for quantifications (**Fig. S3a, c, Fig. S4a, c**). To compare the strain gradients, plots were linearly fitted to acquire slopes (**Fig. S3b, d, Fig. S4b, d**). Three slopes were obtained for reference lines L1 to L3 for each sample. One-way ANOVA test was used to compare individual samples using the slopes for both designs. No significant differences were found.

### Microscopy for cell migration trajectory

An epifluorescence microscope (Leica, DMi8) was used to image the cell migration trajectories. During the experiment, the device was kept in the 37 °C incubators for cell culture. We placed the device onto the microscope stage outside the incubator for imaging, which would take less than 2 mins each time for each sample in the cell migration tracking experiment using the 10× phase contrast. Small air bubbles might be trapped underneath the corner of the cell culture area, which could be removed by lightly knocking the device one or two times. For the consistency of the culture condition, all samples would be subjected to the same knocking motion before imaging. The images were acquired every 50 mins for the static strain condition and every 30 mins for the cyclic strain condition for 6 hrs.

The obtained image sequences were aligned by tracking two reference points that stayed at the constant location on the substrate for the entire experiment period. The reference points could be small particles left on the sample surface. The first image acquired after the stretching step was set to be the fixed image. We used the Matlab *fitgeotrans* function for the translation and the *imrotate* function for the rotation to align the image sequence, allowing clear cell migration trajectories for tracking. Before cell tracking, we would draw lines on the image sequences to encircle the region of interest (**Fig. 2a**). All cells with clear migration trajectories within the encircled region were tracked with the ImageJ MTrackJ plugin, and the direction angles were quantified using customized Matlab codes. The representative cell tracking movies for five experiment groups were shown in **Video S1-S5**.

### Device modification for focal adhesion imaging

We adjusted the designs of the device to allow high-resolution imaging for focal adhesion analysis. To minimize the distance between the cells and the objective, we specifically designed a glass bottom PDMS dish (**Fig. S8.b**). 30 g of PDMS was poured into a 60 mm Petri dish to form a dish-shaped PDMS slab. A 20×46 mm cuboid was cut off in the middle, and then the opening was covered by plasma-bond a thin 24×50 mm cover slide. The dish was baked in a 65 °C oven overnight for firm bonding. The removable bottom of the device was redesigned as well (**Fig. S1**). We laser cut a 60 mm diameter opening on the removable acrylic bottom. The opening was designed to hold the PDMS dish properly, and small gaps were left for manual adjustment of the height of the PDMS dish during focal adhesion imaging.

To mitigate the interference of the PDMS membrane in the light path, we flipped the double-layer membrane to make cells face down during imaging (**Fig. S8**). To fit the cell culture sample into the PDMS dish, the glass slide was cut into smaller 20×14 mm pieces (**Fig. S8b**). The PDMS membrane and the silicone film base were bonded at the first step. The sample was then flipped to bond the glass slides on the opposite side of the PDMS membrane (**Fig. S8a**). As a result, when the adjusted double-layer membrane was mounted onto the device, the length of the silicone film was the same as the regular triangular sample, making the cells exposed to the same strain gradient and stretch magnitude as the cell tracking experiments. The sample was baked at 65 °C in an oven overnight for firm bonding after both steps, and then mounted onto the device. During the experiment, we flipped the device to coat FN and seeded cells on the bottom side of the PDMS membrane. Then the device was put into the incubator upside down for cells to attach for 1 hour. We then flipped the device back and added 3 to 5ml culture media into the PDMS dish. The culture media component was adjusted by replacing the DMEM with the non-phenol red type (21063029, Gibco). Other reagents remained the same.

Right after taking the first image, the cells were stretched once, and then the device was placed back into the incubator for 10 mins before taking the second image. The time gap between two focal adhesion images was within 20 mins. A 40× objective and YFP cube were used to image the focal adhesions.

### Focal adhesion and protrusion/retraction image processing and quantification

The focal adhesion raw images were processed through ImageJ. Briefly, background was removed with the *Subtracted Background* function using the *Sliding paraboloid* option. To enhance contrast, the *CLAHE* plugin was used with the *Blocksize* set to 19, *Histogram bins* set to 256, and the *Maximum slope* set to 6 (50). We further adjusted the *Brightness/Contrast* automatically to increase brightness. The focal adhesions were enhanced at this point. We turned the image format into 8 bits and adjusted the threshold automatically. The sizes and the coordinates of focal adhesions were quantified with the *Analysis particle* function.

The fluorescence intensity declined after cell stretching. Therefore, to effectively compare the focal adhesion dynamics within each cell, the data was normalized using images prior to stretching. The normalization was calculated as below:

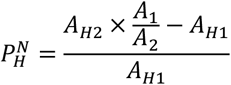

where *A*_1,_ *A*_*H*1_ are the focal adhesion size for the entire and one half of the cell before stretching, respectively; *A*_2,_ *A*_*H*2_ are the focal adhesion size for the entire and one half of the cell after stretching, respectively; and *P*_*HN*_ is the normalized percentage change of focal adhesion sizes in the respective half of the cell.

To quantify protrusion and retraction, we manually encircled the cell and the nuclei boundaries with dots using the ImageJ *MTrackJ* plugin. (**Fig 4. a, c**). The dots were connected automatically with Matlab to generate the boundaries. For each cell, the images before and after stretching were aligned with the mass center of the nucleus and the maximum gradient direction axis. The cell spreading areas were automatically quantified using the *Analysis particle* function.

### Implementation of the extended motor-clutch model

We develop an EMM to simulate the interactions of cells and the elastic substrate based on the previously established model (42-43). Briefly, the cell was simplified as a polygon, with the vertexes connecting to the elastic substrate with integrin-FN bonds. The randomly engaged bonds transmitted cell traction forces between cells and substrates. The on-rate and off-rate of the bonds control stochastic cell-substrate interactions. We assume that the vertex exposed to lower strain would be larger since it would naturally bind the substrate easier. As such, the on-rate *k*_*on*_ would be larger at the lower strain side. Based on a previous study (51), we propose that the on-rate and off-rate of integrin-FN bonds can be calculated as

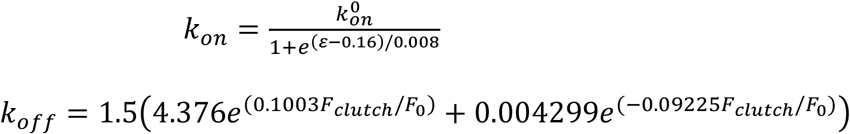

in which 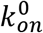 and *ε* represent the reference on-rate and the local strain of substrates, respectively. *F*_*clutch*_ = *k*_*c*_(*x*_*c*_ − *x*_*sub*_) is the clutch force of a single integrin–FN bond. *k*_*c*_ is the spring constant, *x*_*c*_ is the displacement of bond on the filament, and *x*_*sub*_ is the substrate displacement. *F*_0_ = 1*pN* is the reference force magnitude to normalize *F*_*clutch*_. With engagement of integrin-fibronectin pairs, the substrate displacement can be calculated from

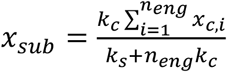

where *n*_*eng*_ is the number of the engaged bonds on the filament and *x*_*c,i*_ is the displacement of the *i*^*th*^ bond on the filament. According to Cerruti problem, spring constant of substrate *k*_*s*_ can be calculated by (42):

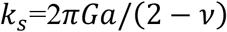

where *v* is the Poisson ratio and *G* = *E*/2(1 + *v*).

The cell migration can be modeled by the motion equation of each vertex, which is given by:

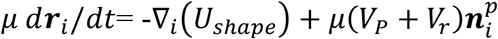

where *μ* is the friction coefficient. The first term is constituted by contraction force 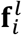, area constraint force 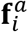 and bending force 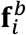 (**Fig.5a**) (42). The second term is the cell polarization force, where *V*_*P*_ and *V*_*r*_ are the polymerization and retrogradation flow speed along the orientation of each filament 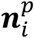, respectively. Retrogradation flow velocity is determined by cell tractions (43) and can be calculated as

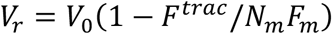

where *V*_0_ is the unloading velocity of each filament and *F*^*trac*^ = *k*_*s*_*x*_*sub*_ is the total traction force magnitude exerted by the filament. *N*_*m*_ is the number of myosin motors, whose stall force is denoted by *F*_*m*_. The myosin motors exert cell contractions along filaments. Our simulations are implemented on Matlab, and detailed parameters are provided in **Table S1**.

### Statistics

Statistical analysis was performed using OriginLab and R. Rayleigh tests were used to determine the unimodal distribution of circular data in cell migration direction analysis (52). For non-circular datasets, normality tests were run to determine the normality of the distribution. For statistical comparisons of two normal distributed datasets, P-values were calculated using the two-sample t-test. For statistical comparisons of two non-normal distributed datasets, P-values were calculated using the non-parametric Mann-Whitney test. For statistical comparisons of multiple normal distributed datasets, P-values were calculated using the one-way ANOVA test.

## ACKNOWLEDGMENT

This work is supported in part by the National Science Foundation (CMMI 1846866 to Y. Sun.) and the National Institute of Diabetes and Digestive and Kidney Diseases (R01DK129990 to Y. Sun).

## AUTHOR CONTRIBUTIONS

F. Y. designed and performed the experiments, analyzed the data, and wrote the manuscript. P. C. performed the simulation and wrote the manuscript. T. X. microfabricated micropatterned stamps. Y. Sun, B. L., and Y. Shao supervised the project. All authors edited and approved the manuscript.

## DECLARATION OF INTERESTS

The authors declare no competing interests.

## Notes

### Competing Interest Statement

The authors have declared no competing interest.

### Summary of Updates

New motor-clutch modeling is added in this version (Fig. 5)

