## Supplemental Figures and Tables for "Directional Cell Migration Guided by a Strain Gradient"

**This PDF file includes:**

Figures S1 to S8
Tables S1 to S2
Legends for Movies S1 to S5
Supplementary References

**Other supplementary materials for this manuscript include the following:**

Movies S1 to S5

**Supplementary Figures**

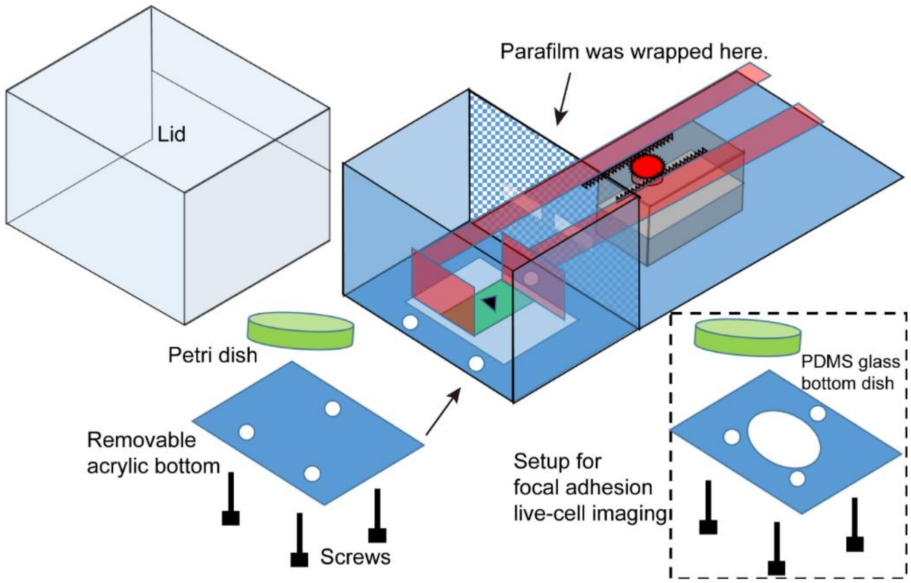


**Fig. S1. Schematics showing the assembly of the cell stretching device.** The removable bottom of the device can either fit a regular 60 mm petri dish for cell migration assay, or a customized glass bottom PDMS dish for imaging focal adhesions.

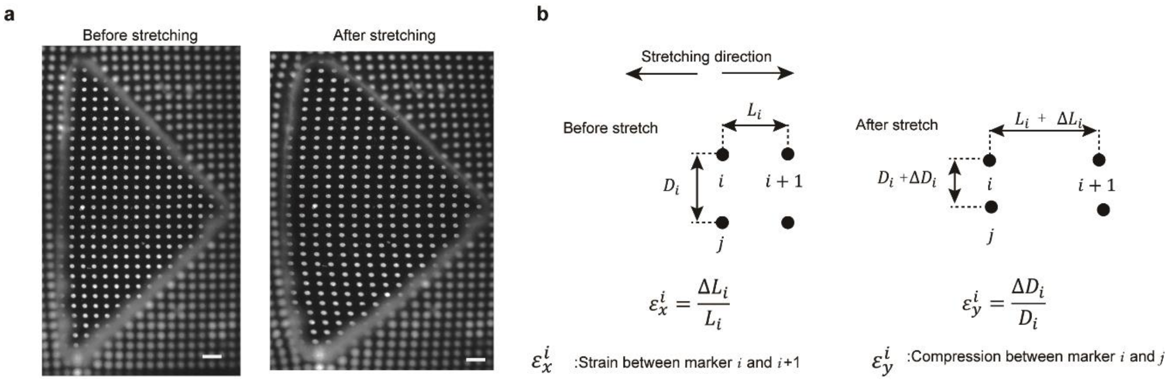


**Fig. S2. Calibration of the strain field.** (a) Fluorescence images showing of membranes with triangle cut-out stamped with fluorescent marks before (left) and after (right) stretching. Scale bar, 200 µm. (b) Schematic showing the method for strain and compression calculation.

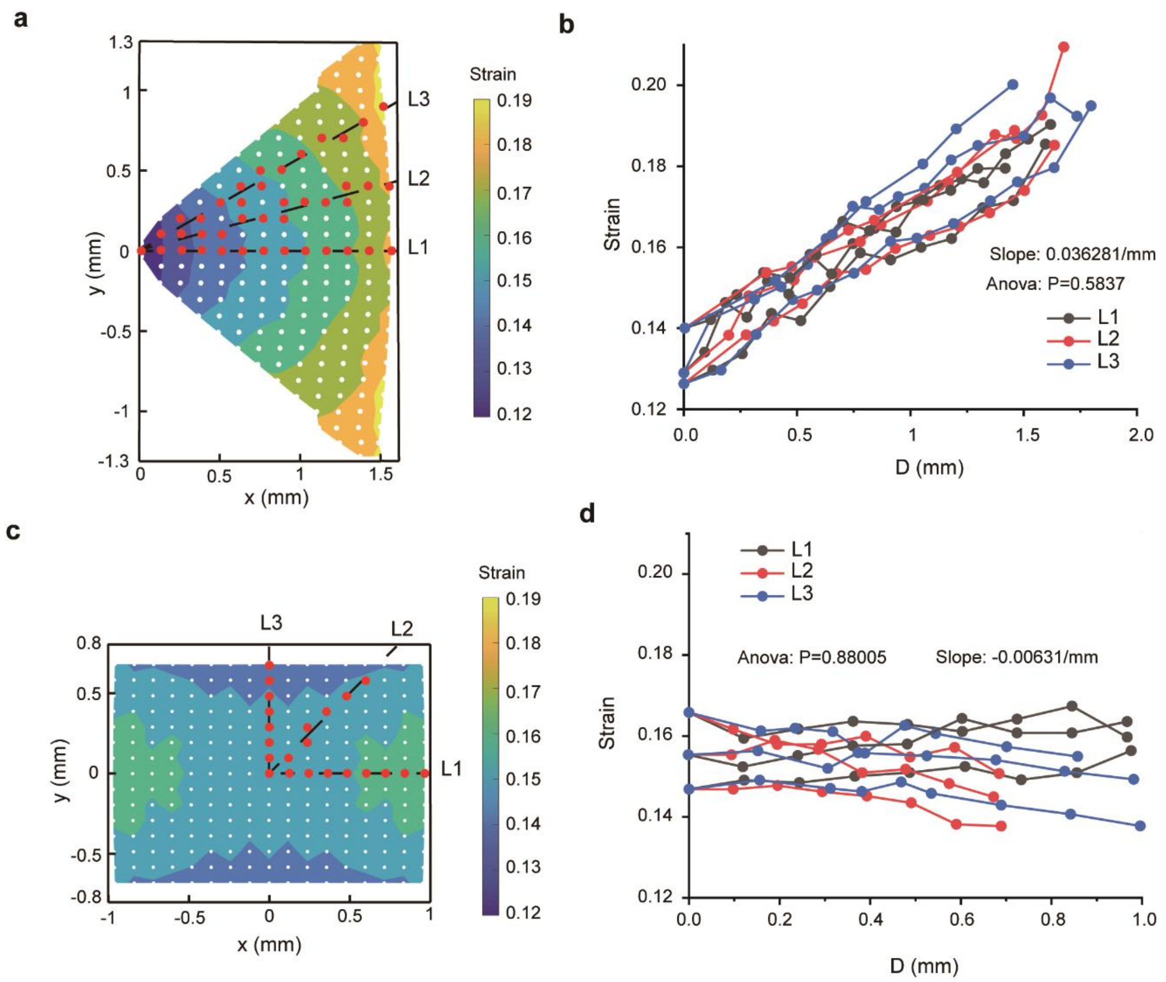


**Fig. S3. Reproducibility of the strain field generated on different samples.** (a) A colorimetric map showing experimentally measured strain field generated on the membrane with triangle cut-out. Both red and white dots showed locations where strains were measured. The strains along three directions (L1, L2, L3; marked in red) as a function of the distance (D) from the triangle's vertex were plotted in (b). (c) A colorimetric map showing experimentally measured strain field generated on the membrane with square cut-out. Both red and white dots showed locations where strains were measured. The strains along three directions (L1, L2, L3; marked in red) as a function of the distance (D) from the center of the square were plotted in (d). ANOVA tests were performed to determine the statistical significance of the variation among individual samples using the slopes of the plots. N = 3 for both designs.

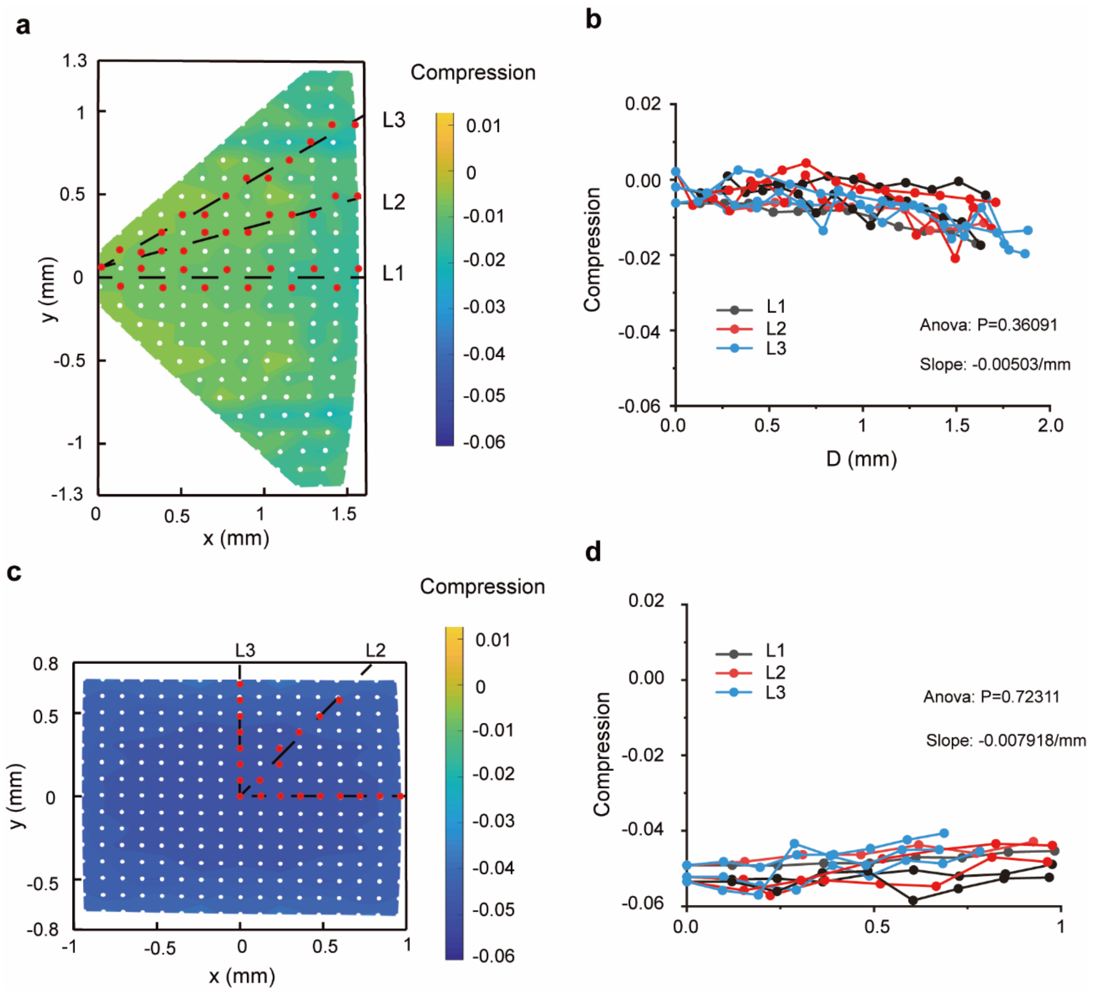


**Fig. S4. Reproducibility of the compression field generated on different samples.** (a) A colorimetric map showing experimentally measured compression field generated on the membrane with triangle cut- out. Both red and white dots showed locations where compressions were measured. The compressions along three directions (L1, L2, L3; marked in red) as a function of the distance (D) from the triangle's vertex were plotted in (b). (c) A colorimetric map showing experimentally measured compression field generated on the membrane with square cut-out. Both red and white dots showed locations where compressions were measured. The compressions along three directions (L1, L2, L3; marked in red) as a function of the distance (D) from the center of the square were plotted in (d). ANOVA tests were performed to determine the statistical significance of the variation among individual samples using the slopes of the plots. N = 3 for both designs.

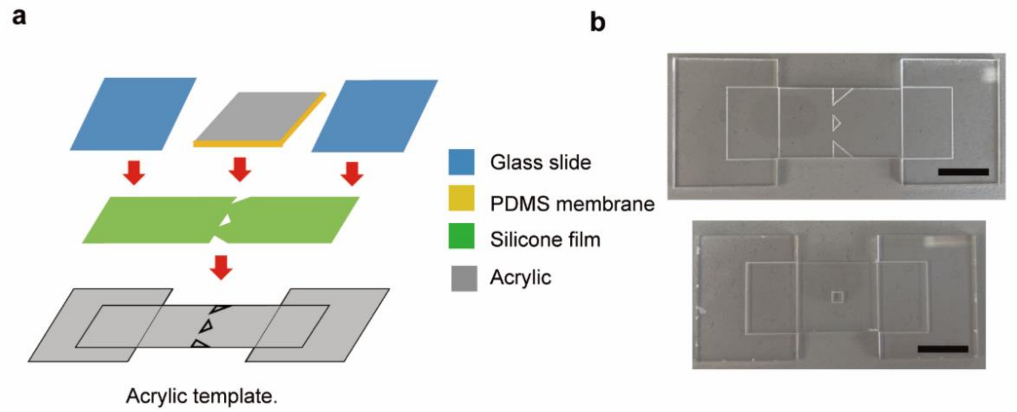


**Fig. S5. Fabrication of the double-layer membrane for cell culture and stretching.** (a) Schematic of the fabrication procedure. (b) Photos of the acrylic templates with alignment marks for samples with triangle (top) and square (bottom) cut-out. Scale bar, 10 mm.

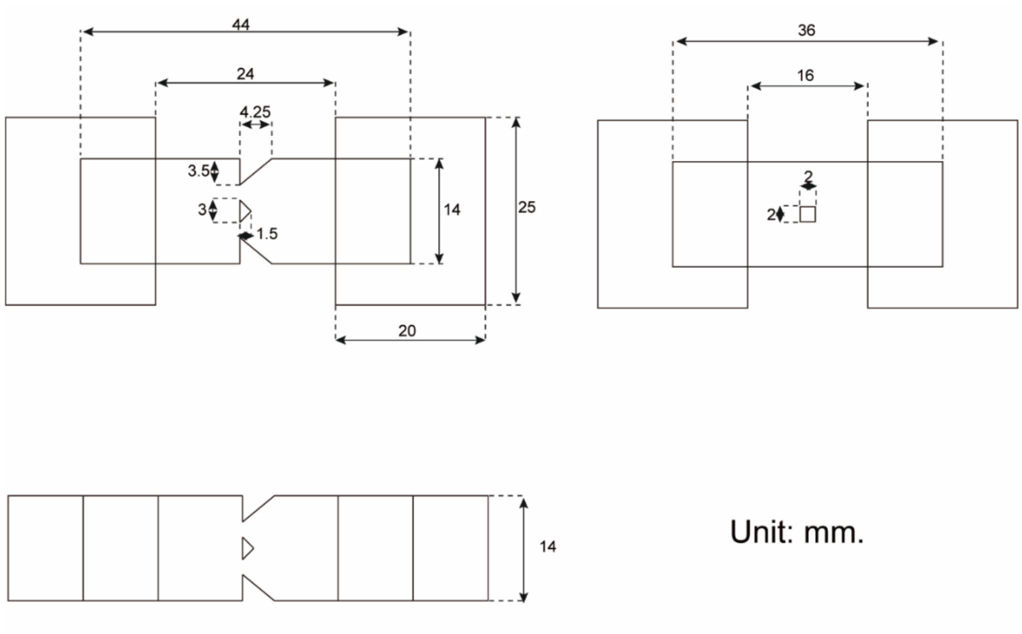


**Fig. S6. The dimensions of double-layer membranes.** Top left: triangle design; Top right: square design; Bottom: adjusted triangle design for focal adhesion imaging.

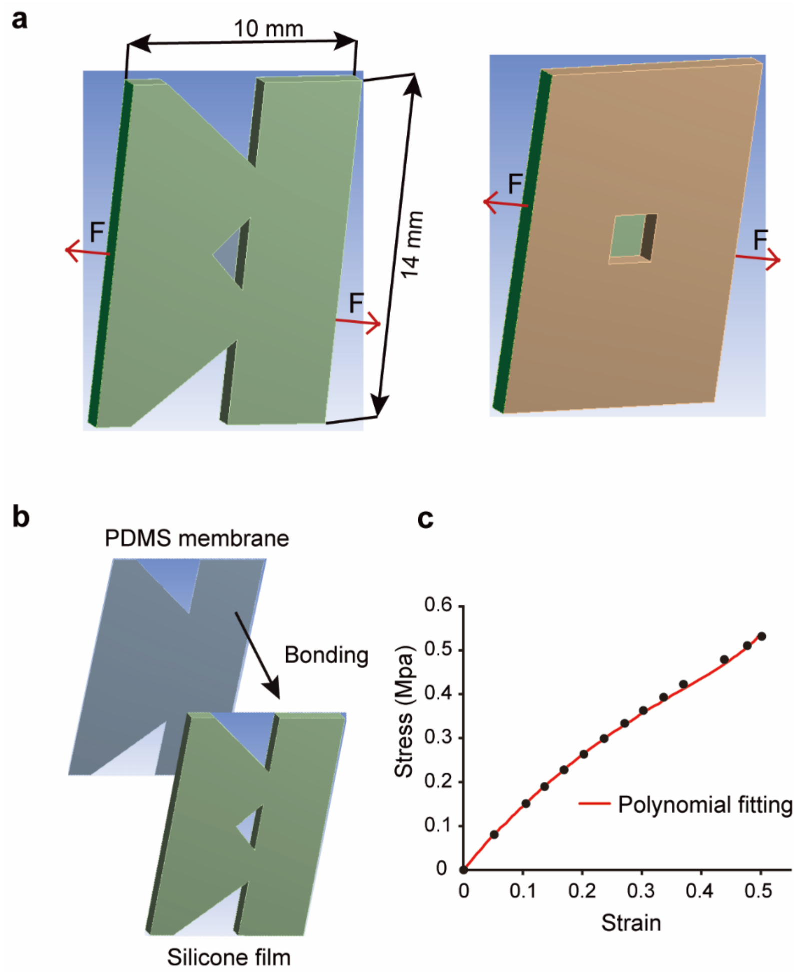


**Fig.S7. Finite element modeling of the double-layer membrane.** (a) 3D models for simulating the silicone membrane with triangle and square cut-out. (b) 3D models showing the bonding of the silicone base to the PDMS membrane. (c) Stress-strain curve of silicone film measured using a force gauge fitted using the polynomial 2nd order equation.

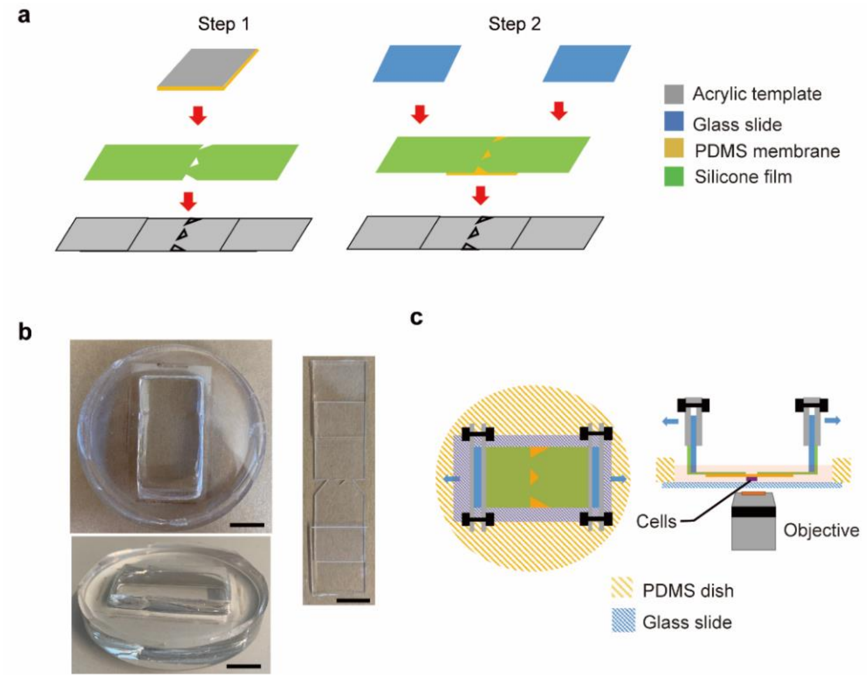


**Fig. S8. Device modification for focal adhesion live-cell imaging.** (a) Schematic of adjusted membrane fabrication procedure. (b) Photos of the glass bottom PDMS dish (left, showing both top view and side view) and the modified membrane (right). Scale bar, 10 mm. (c) Schematic of imaging and sample stretching of the modified sample.

**Supplementary Tables**

**Table S1. Parameters for the Extended Motor-clutch simulation**

| Parameter | Symbol | Value | Ref. |
|---|---|---|---|
| Initial cell radius | $R$ | 25 μm | |
| Number of vertexes | $N_v$ | 34 | 1 |
| Target edge length | $l_0$ | $2Rsin(\pi/N_v)$ | |
| Target area | $a_0$ | $R^2 sin(2\pi/N_v)/2$ | |
| Damping coefficient | $\mu$ | 0.005 Ns/m | 2 |
| Contractive modulus | $k_l$ | $10^{-4}$ N/m | 2 |
| Area modulus | $k_a$ | $4.34 \times 10^6$ $Nm^{-3}$ | 2 |
| Bending energy | $k_B$ | $10^{-14}$ J | Assumed |
| Number of myosin motors | $N_m$ | 135 | 3 |
| Stall force of a single myosin motor | $F_m$ | -2 pN | 4 |
| Retrogradation speed of unloaded filament | $V_0$ | -110nm/s | 3 |
| Polymerization speed of actin filaments | $V_P$ | 120nm/s | 5 |
| Number of clutches | $N_f$ | 75 | 3 |
| Factor of binding rate | $k_{on}^0$ | 0.3/s | 6 |
| Elastic coefficient of clutches | $k_c$ | 1 nN/nm | 7 |
| Stiffness of substrate | $E$ | 1.1MPa | 8 |
| Poisson's ratio | $\nu$ | 0.5 | Assumed |
| Time interval | $\tau$ | 5ms | |
| Simulation time | $T$ | 6 h | Assumed |

**Table S2. Parameters used in the finite element modeling.**

| | |
|---|---|
| PDMS membrane | Thickness: 0.1 mm Young's modulus: 1.1Mpa Poisson's ratio: 0.45<br>Density: 970 kg/$m^3$ |
| Silicone base | Thickness: 0.79 mm |
| Mesh size | 0.15 mm |

| | |
|---|---|
| Force in x direction | Triangle design: F = ±0.42 N<br>Squire design: F = ±0.47 N |

**Movie S1 (separate file).** Representative video showing tracked cell migration under a static strain gradient.

**Movie S2 (separate file).** Representative video showing tracked cell migration under a static uniform strain.

**Movie S3 (separate file).** Representative video showing tracked cell migration when the membrane was pre-stretched under a static strain gradient before cell seeding.

**Movie S4 (separate file).** Representative video showing tracked cell migration under a cyclic strain gradient.

**Movie S5 (separate file).** Representative video showing tracked cell migration under a cyclic uniform strain.
